# Cethromycin Pharmacokinetics and Pharmacodynamics for Single Dose Cure of *Plasmodium berghei* Liver Stages

**DOI:** 10.1101/2025.02.10.637401

**Authors:** Grace Kennedy, Rachel M. West, Kristin Poti, Bryce Bobb, Matthew M. Ippolito, Mark A. Marzinke, Nikola Kaludov, David J. Sullivan

**Author notes:** Corresponding author: David J. Sullivan, Johns Hopkins Bloomberg School of Public Health. 615 North Wolfe Street, Baltimore, Maryland 21205, USA.

## Abstract

Cethromycin combines a quinoline nucleus and a macrolide for broad spectrum antibacterial and antiprotozoan activity. Here we characterized the murine pharmacokinetics and *Plasmodium berghei* lifecycle stage pharmacodynamics for the cethromycin base. Liver pharmacokinetic studies in mice show peak mM drug levels in the liver with 20 hour sustained levels above 10 μM. Peak concentrations in the liver were double the lung and about 440 times that of plasma. Immunofluorescence imaging of *in vitro* cethromycin-treated infected hepatocytes shows complete ablation of the apicoplast. We observed complete cure of *P. berghei* liver stage infection by single oral dose of 60 mg/kg in mice which is equivalent to the 5 mg/kg human dose of 300 mg a day used in bacterial pneumonia studies. Cethromycin at 60 mg/kg daily for 7 days was curative in the high parasitemic *P. berghei* mouse model. Both mosquito membrane feeding of *P. falciparum* gametocytes incubated with 20 μM cethromycin and oral dosing in mice demonstrated no decrease in oocyst numbers. Cethromycin has been evaluated for efficacy against bacterial pneumonia in more than 5,000 patients with good safety profiles. Cethromycin has potential for rapid clinical development for casual malaria prophylaxis and possibly radical cure of dormant liver *P. vivax*.

## INTRODUCTION

Cethromycin is a ketolide class macrolide (erythromycin base with a single sugar and quinoline unsubstituted nucleus replacing the cladinose saccharide) with broad spectrum antibacterial, mycobacterial and antiprotozoal activity developed for community acquired pneumonia(1). The drug was safe, but not superior to clarithromycin in a large phase 3 outpatient study of more than 1000 participants(2) for which the new drug application was later denied for the nonsuperior efficacy results by the FDA(1, 3). Cethromycin binds to bacterial 23S ribosomal RNA of the 50S ribosome subunit which stops polypeptide amino-acid addition, terminating protein synthesis and growth(4). Cethromycin specifically binds domains II and V in the wall of the polypeptide exit tunnel(1). While ketolides like cethromycin have greater activity than azithromycin against some bacterial species(5),both are unable to overcome high level *erm* resistance(6–8).

The macrolide class has demonstrated delayed death activity *in vitro* and *in vivo* against protozoan apicomplexans like *Toxoplasma*(9) and *Plasmodium*(*10–12*). The plant-like apicoplast organelle is essential in production of isoprenoids for blood stage parasites. Previous work has shown that isoprenoid supplementation with azithromycin treatment results in loss of the extrachromosomal apicoplast genome and of the organelle in erythrocyte stages(13).

During the liver stages of malaria infection, the macrolide molecule has been shown to also delete the apicoplast and the extrachromosomal apicoplast genome(14). Thus, while the liver stages of malaria infection progress, loss of the apicoplast prevents infection of erythrocytes, thus transition to the blood stage (15). Azithromycin and doxycycline can cure mouse liver stages(16), but in controlled human malaria challenge studies, only 4 of 10 patients treated with azithromycin(17) and 8 of 12 patients treated with doxycycline (18)were cured after 7 days of liver stage treatment initiated on sporozoite challenge.

In a previous report, we described liver-stage *Plasmodium* inhibition in the quantitative PCR mouse liver-stage sporozoites-challenge model with single oral low doses of cethromycin at 15, 25 and 50 mg/kg, because of access to only a few milligrams of newly synthesized cethromycin(19). Here we report more extensive dose-ranging and schedules for cethromycin pharmacokinetics and pharmacodynamics in a murine malaria liver, blood and gametocyte stage models.

## Results

### Pharmacokinetics

We compared cethromycin concentrations in liver, lung and plasma with single oral doses in mice at 30, 60 or 120 mg/kg, which is allometrically equivalent to 150, 300 and 600 mg total oral dose in humans. The maximum concentration (C_max_) in the liver was double that of the lung and approximately 440 times more than plasma. The 765 gm/mole cethromycin molecular weight indicates 1 μM levels at 0.76 mcg/mL or gram in the liver. The time above 1 μM in the liver was 24 hours for the 60 mg/kg dose. Plasma concentrations never exceeded 10 μM and stayed above 1 μM for 12 hrs for the 60 mg/kg dose which is allometrically equal to 300 mg single oral dose in humans (Figure 1). The tissue disposition of cethromycin measured as C_max_, area under the curve (AUC_0-72_), and elimination half-life (t_1/2α_) are shown in Table 1.

**Fig 1.**
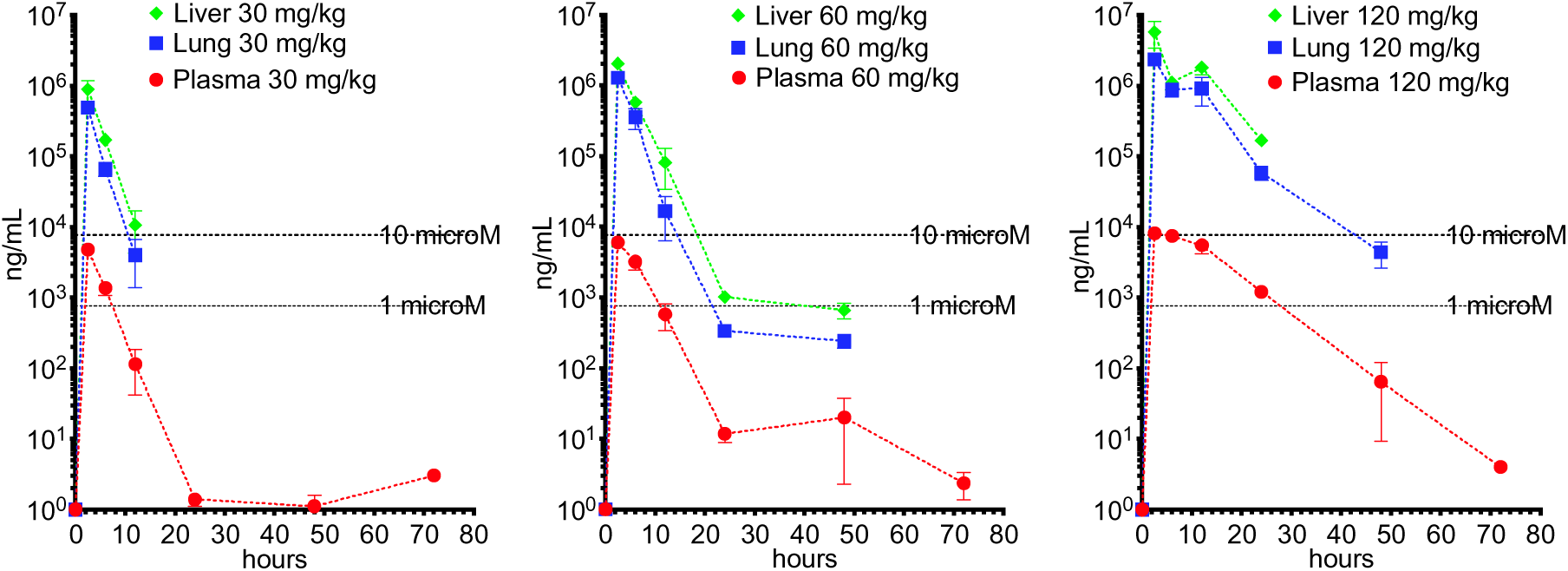
Liver, lung blood pharmacokinetics-. Single oral cethromycin doses were given to groups of three mice with anticoagulated blood, liver and lung harvested, and frozen on dry ice after separation of plasma. A) 30 mg/kg B) 60 mg/kg C) 120 mg/kg. Graphs are mean and standard deviation. Dotted lines are 1 and 10 μM at 765 and 7650 ng/mL respectively

**Table 1.** Tissue distribution of cethromycin after single oral doses in mice (n=3 per group)

| Tissue | Dose (mg/kg) | C<sub>max</sub> (ng/ml) | AUC<sub>0-72</sub> (ng·h /ml) | t<sub>1/2α</sub> (h) |
| --- | --- | --- | --- | --- |
| Plasma | 30 | 4,767 ± 754 | 18,682 ± 2,216 | 1.77 (1.56-2.03) |
| Plasma | 60 | 5,994 ± 700 | 34,429 ± 3,572 | 2.32 (2.40-2.69) |
| Plasma | 120 | 8,297 ± 604 | 117,134 ± 9,894 | 7.53 (5.82-10.66) |
| Lung | 30 | 481,883 ± 45,681 | 1,502,310 ± 122,398 | 1.90 (1.58-2.39) |
| Lung | 60 | 1,291,870 ± 189,700 | 4,856,050 ± 616,430 | 1.70 (1.48-2.00) |
| Lung | 120 | 2,322,030 ± 83,339 | 17,565,300 ± 2,061,010 | 4.21 (3.31-5.80) |
| Liver | 30 | 1,098,800 ± 253,711 | 3,585,590 ± 596,799 | 1.85 (1.61-2.17) |
| Liver | 60 | 2,063,840 ± 74,153 | 8,573,030 ± 451,875 | 2.02 (1.85-2.23) |
| Liver | 120 | 6,221,820 ± 1,501,440 | 37,067,800 ± 4,292,660 | 4.73 (3.64-6.74) |

Mean and s.d. maximum concentration (C<sub>max</sub>) and area-under-the-curve (AUC<sub>0-72</sub>) are shown. The 24-hour half-life (t<sub>1/2α</sub>) was estimated from linear regression of the log- transformed data with 95% confidence intervals shown in parentheses.

### Liver stage apicoplast disruption

We investigated maturation of the liver stage *P. berghei* in vitro in 48-hour hepatocyte cultures treated with primaquine, azithromycin and cethromycin exposure at 20 mM resulted in parasites with predominately no observed apicoplasts. Detailed counting of infected hepatocytes noted that those exposed to primaquine had parasites with no apicoplast observed in all hepatocytes, but single dysmorphic apicoplasts were present in parasites in cethromycin and azithromycin treated hepatocytes. In the controls, all apicoplasts were highly branched indicating normal hepatocyte development of the parasite. The heat shock protein (hsp) 70 reporter also indicated almost normal parasite development with azithromycin and cethromycin exposure nearly indistinguishable from control while primaquine-exposed parasites contained small pyknotic *P. berghei* hsp 70 signal (Fig 2).

**Fig 2.**
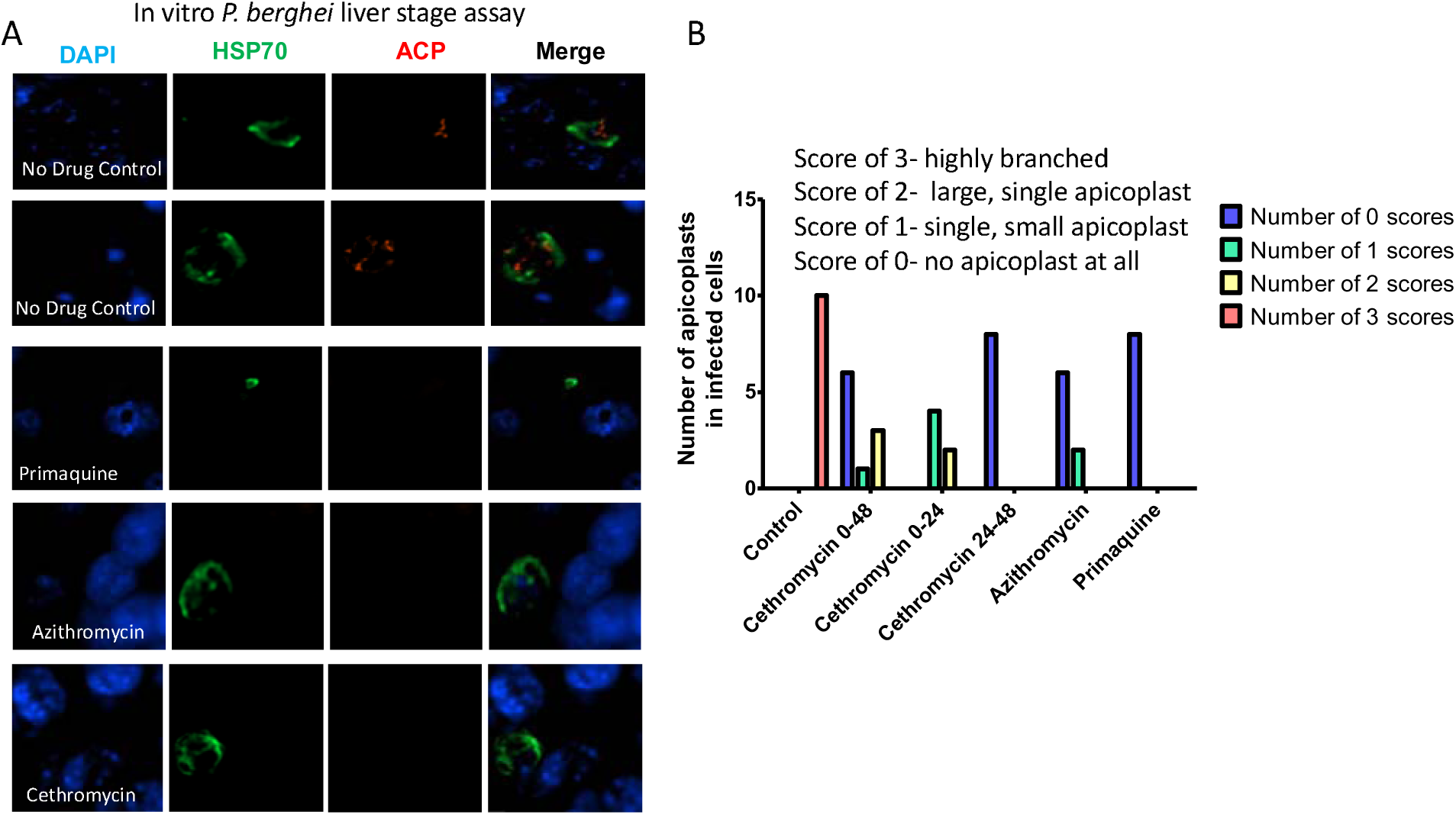
Hepatocyte *Plasmodium berghei* apicoplast imaging after drug treatment-. Drugs at 20 μM were incubated 2 hours after sporozoite Hepa-1 hepatocyte invasion for 48 hours at which time cells were fixed for immunofluorescence. A) Representative immunofluorescent images at 48 hours stained for nuclei with DAPI, cytosol with *P. berghei* heat shock protein 70 (HSP70) and the apicoplast with acyl carrier protein (ACP). B) Segregation of number size and branching of the apicoplast in hepatocytes under experimental conditions with the addition of cethromycin incubated for the first 24 hours and last 24 hours in addition to 48 hour drug incubations. Experiments were done in biologic replicates.

### Liver stage pharmacodynamics

In the 3000 sporozoite tail vein injection murine liver-stage model repeated on three separate occasions, all 9 controls had patent bloodstream parasites, while none of the mice treated with cethromycin at 60 mg/kg single oral dose 24 hours after sporozoite injection were patent. All 6 mice dosed with single oral 15 mg/kg cethromycin were patent and 1 of 2 at the 30 mg/kg dose. Utilizing a more natural lower dose of sporozoites with 5-8 infected mosquito bites, all controls were positive in the 8 separate biologic dosing experiments shown in Table 2 and Fig 3. Cethromycin dosed at 30 or 60 mg/kg 24 hours prior to infection cured zero mice. Both azithromycin and cethromycin dosed 2 hours after mosquito infection were curative at 60 mg/kg oral dose. Doses 24 hours after infection demonstrated 47% patency with azithromycin when combining results of the 30 mg/kg (3/6 patent), 45 mg/kg (1/3 patent) and 60 mg/kg (3/6 patent) doses. For cethromycin 30 mg/kg (1/9 patent), 45 mg/kg (0/3 patent) and 60 mg/kg (1/9 patent) more mice were cured. Cethromycin 20 mg/kg dosed at day -1, 0 and 1 was curative in all mice (n=3). Quinoline nucleus and erythromycin ethyl succinate, the two components of cethromycin, given alone and in combination were not curative when dosed on day 0 and 1 at 120 mg/kg. C57Bl/6 mice are more permissive/susceptible to murine malaria liver stages than Balb/c with higher liver loads given the same inoculum. In C57Bl/6 mice azithromycin 45 mg/kg had one patent mouse while at same dose of cethromycin 0 of three were patent. At the 60 mg/kg dose all nine mice were cured.

**Table 2.** Liver stage patency.

| Mouse | # Mosq | Drug Groups | Dose mg/kg | day of dose | number pos/total | average day to patency | number positive | total number |
| --- | --- | --- | --- | --- | --- | --- | --- | --- |
| BALB/c | 7 | Cethromycin | 30 | -1 | 3/3 |  |  |  |
| BALB/c | 7 | Cethromycin | 60 | -1 | 3/3 |  |  |  |
| BALB/c | 7 | Azithromycin | 45 | 0 | 3/3 | 8 | 3 | 3 |
| BALB/c | 7 | Azithromycin | 60 | 0 | 0/6 |  | 0 | 6 |
| BALB/c | 7 | Azithromycin | 120 | 0 | 0/3 |  | 0 | 3 |
| BALB/c | 7 | Cethromycin | 45 | 0 | 1/3 | 8 | 1 | 3 |
| BALB/c | 7 | Cethromycin | 60 | 0 | 0/3 |  | 0 | 3 |
| BALB/c | 7 | Cethromycin | 120 | 0 | 0/2 |  | 0 | 2 |
| BALB/c | 7 | Azithromycin | 15 | 1 | 2/3 | 7 | 2 | 3 |
| BALB/c | 7 | Azithromycin | 30 | 1 | 3/6 | 9 | 3 | 6 |
| BALB/c | 7 | Azithromycin | 45 | 1 | 1/3 | 13 | 1 | 3 |
| BALB/c | 5 | Azithromycin | 60 | 1 | 3/6 | 9 | 3 | 6 |
| BALB/c | 7 | Cethromycin | 10 | 1 | 3/3 |  |  |  |
| BALB/c | 7 | Cethromycin | 15 | 1 | 5/12 | 8 | 5 | 12 |
| BALB/c | 7 | Cethromycin | 30 | 1 | 1/9 | 10 | 1 | 9 |
| BALB/c | 7 | Cethromycin | 45 | 1 | 0/3 |  | 0 | 3 |
| BALB/c | 5 | Cethromycin | 60 | 1 | 1/9 | 13 | 1 | 9 |
| BALB/c | 5 | Cethromycin | 120 | 1 | 0/3 |  | 0 | 3 |
| BALB/c | 5 | Primaquine | 40 | 1 | 0/3 |  | 0 | 3 |
| BALB/c | 5 | Primaquine | 60 | 1 | 0/3 |  | 0 | 3 |
| BALB/c | 7 | Cethromycin | 60 | 2 | 3/3 | 6 |  |  |
| BALB/c | 7 | Cethromycin | 60 | 3 | 3/3 | 6 |  |  |
| BALB/c | 7 | Cethromycin | 20 | -1, 0, 1 | 0/3 |  |  |  |
| BALB/c | 5 | Cethromycin | 120 | 0,1 | 0/3 |  | 0 | 3 |
| BALB/c | 5 | Erythromycin Ethyl Succinate | 120 | 0,1 | 2/2 | 6 | 2 | 2 |
| BALB/c | 5 | Quinoline | 120 | 0,1 | 3/3 | 4 | 3 | 3 |
| BALB/c | 5 | Primaquine | 60 | 1,1 | 0/3 |  | 0 | 3 |
| BALB/c | 7 | Untreated |  |  | 24/24 | 6 | 24 | 24 |
| C57Bl6 | 7 | Azithromycin | 45 | 1 | 1/3 | 13 | 1 | 3 |
| C57Bl6 | 7 | Cethromycin | 45 | 1 | 0/3 |  | 0 | 3 |
| C57Bl6 | 8 | Cethromycin | 60 | 1 | 0/9 |  | 0 | 9 |
| C57Bl6 | 8 | Untreated | n/a |  | 3/3 | 5 | 3 | 3 |

**Fig 3.**
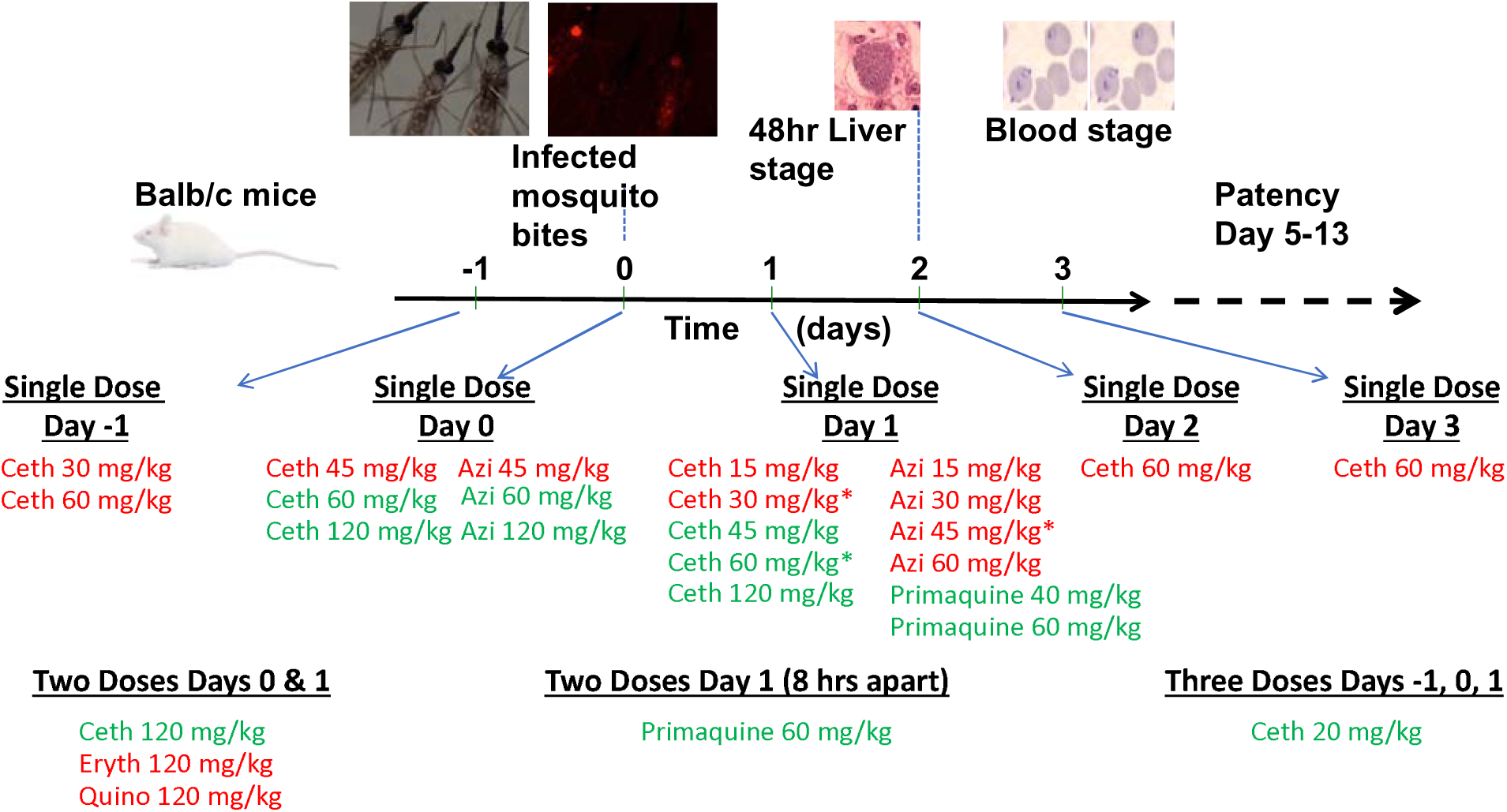
Schema for liver stage dosing schedule with drugs dosed before during and after 48 hours *P. berghei* liver stage. Infected mosquitos were sorted with fluorescent microscopy for mcherry sporozoites and 5-7 mosquitoes were fed on anesthetized mice. Patency was measured by Giemsa blood film for 16 day after sporozoite infection. Red labeled drugs indicate partial or all mice patent and green labeled drugs indicate zero patent mice after liver stage infection.

### Blood-stage

We measured the efficacy of cethromycin and clindamycin in *P. falciparum* isolates 3D7. The 72 hour IC_50_s were in the low mM range while the 144 hour assays indicated 100-fold more potency to near 60 nM IC_50_ (Fig 4A). In the murine *in vivo* 4 day suppression test, drug doses started after a low (one million) parasite intraperitoneal inoculum. Both azithromycin and cethromycin at 60 mg/kg showed a similar >99% inhibition (more than 3 log difference in parasite number) at day 4 but both subsequently rebounded to one million parasites per μL on day 11 post treatment (Fig 4B). The high parasitemia model with 1 million parasites per μL azithromycin showed greater parasite reduction than cethromycin at 2 and 4 days of oral drugs (Fig 4C). In the same high parasitemia model 6 days of cethromycin was partially curative (2/3 mice), while 7 days cured all mice (Fig 4D).

**Fig 4.**
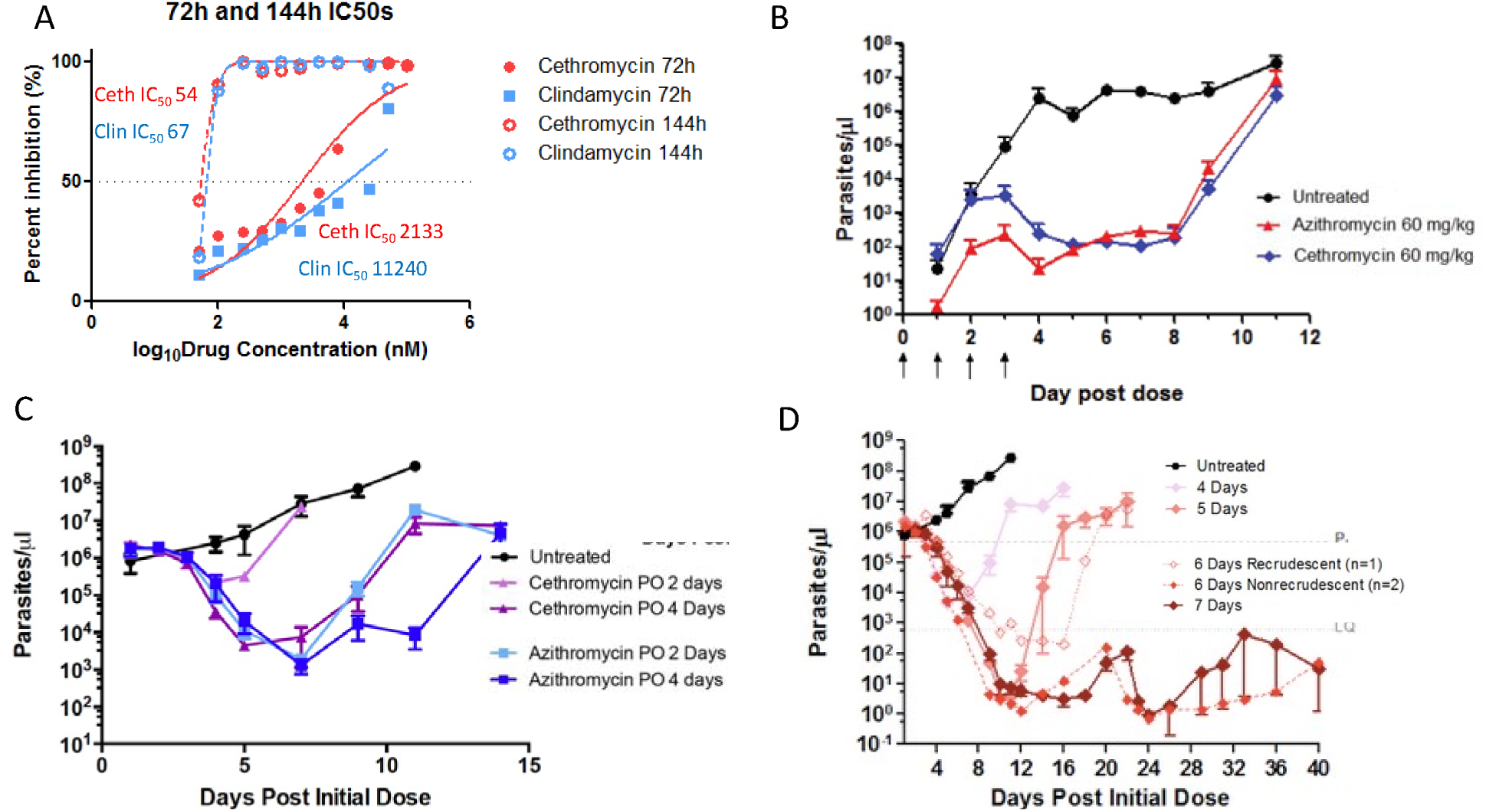
Blood stage inhibition. A) *P. falciparum* inhibition-Continuous 72 and 144 hour *P. falciparum* 3D7 drug incubations with cethromycin and clindamycin. Assays performed in biologic duplicates with technical triplicates. B) *P. berghei* blood stage 4 day suppression test with drugs starting 24 hours after inoculation for 4 days. C) High parasitemia cytocidal test with 2 and 4 days of oral cethromycin and azithromycin D) High parasitemia test model with 4, 5, 6 and 7 days of cethromycin. One of three mice recrudesced with 6 days of dosing and none of the 7 day dosing Female Balb/cj mice (n=3) were inoculated with approximately 1 million infected erythrocytes from a donor mouse. 4-5 days before drug dosing at about 1 million parasites per μL. Data is represented as mean ± SEM.

### Gametocyte stages

Investigation of cethromycin and azithromycin activity against late stage IV and V gametocytes where 15 day old gametocyte cultures were incubated for 48 hours with 10 μM drugs followed by transmembrane feeding noted similar oocyst numbers and infection rates (Fig 5).

**Fig 5.**
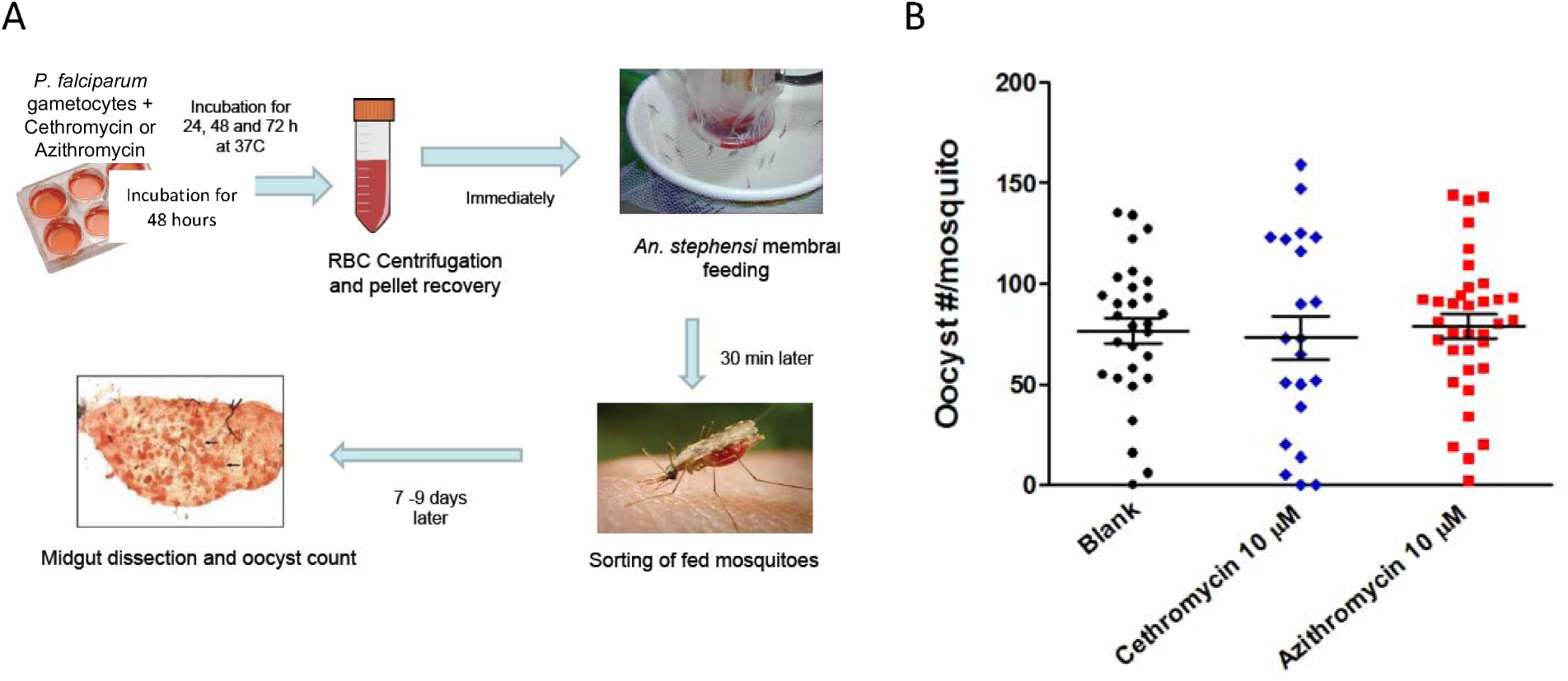
Late stage gametocyte to oocyst test-. A) Schema for late stage IV and V gametocytes on day 15 were dose with 10 μM drug for 48 hours, cells washed and immediately placed in membrane feeding apparatus, after which fed mosquitoes were sorted and number of oocysts per mosquito counted. At least 20 mosquitoes were counted.

## DISCUSSION

In this work, we show that cethromycin demonstrates potent causal prophylactic activity against *P. berghei* malaria liver stages with superior cure rates compared to azithromycin at equivalent doses, while exhibiting typical macrolide delayed-death activity in blood stages and no significant effect on late-stage gametocytes. This activity profile aligns with the known mechanism of macrolide antimalarials, where the inability of the apicoplast to successfully replicate results in parasite death in the second generation of parasites and leads to the phenomenon of delayed generational death(20). Macrolides target both the *Toxoplasma* and *Plasmodium* apicoplast where the tissue stages appear to progress normally, but the progeny all are nonviable and lack functional apicoplasts for isoprenoid production(13). Cethromycin cures liver stage malaria in mice with a single low dose of 60 mg/kg (human equivalent dose of 300 mg). A dose of 20 mg/kg in mice over three days, which is equivalent to a 100 mg single dose in humans, was also curative. Importantly, the two molecular constituents of cethromycin—erythomycin and quinoline—when given alone or together as separate molecules, did not cure any mice. We compared the same dose of a related macrolide in widespread clinical use, azithromycin at 60 mg/kg, and this was not curative. Primaquine was curative at the single dose of 40 mg/kg. The single dose of cethromycin given on day 2 or 3 was not curative, showing that its primary effect is on the liver stage and not subsequent blood stages.

Cethromycin 60 mg/kg, which is allometrically equivalent to the 300 mg single daily dose in humans, achieved maximum liver concentrations of nearly 3 mM and remained above 1 μM for 24 hours(21). Azithromycin liver concentrations dosed at 200 mg/kg were 36 mcg/mL (48 mM) a day after dosing(22), and were 200 times that of serum. Another publication noted a peak liver azithromycin concentration of 73 mcg/mL (97 mM) after 50 mg/kg dose with four times higher concentrations in liver than lung (23). Here we noted 2,063 mcg/mL as the observed peak liver concentration after the 60 mg/kg oral dose, which was measured 2.5 hours post-dosing. These data support other findings which show greatest cethromycin deposition in the liver. Macrolides are ideally suited by pharmacokinetics to work on liver infections.

In separate experiments initiated by blood stage infection and bypassing the liver, 7 daily oral doses were curative of a high blood stage parasite density. This is comparable to the seven days necessary to cure with artesunate—a first-line agent for severe malaria— in the murine mouse model(24). However, the slow parasiticidal activity of macrolides which kill by delayed generational death precludes their use as monotherapy for severe malaria, where rapid parasite reduction is necessary(25). They do have near equal curative activity in the mouse model to artesunate when dosed on a daily basis. Artesunate is superior in rapid parasite clearance activity in humans compared to the antibacterial malaria drugs(26).

The demonstrated cure and an excellent therapeutic product profile of the 3-keto macrolide quinoline cethromycin suggest further investigation into the niche area of dormant liver stage malaria. Recent work noted disruption of the apicoplast in both dormant *P. vivax* and the primate equivalent species *P. cynomolgi*(14) with the speculation that azithromycin might work in relapsing malaria. Azithromycin in a triangular human clinical test design of 5 patients was not active in prevention of dormant *P. vivax* with 3 relapses stopping the trial (27). In the *P. cynomolgi* model clindamycin by itself or with subcurative primaquine did not stop relapse but azithromycin with subcurative primaquine prevented relapse in one of two monkeys, but could not be distinguished from radical cure with primaquine alone in this model(28).

We hypothesize that the quinoline ring might give more liver stage killing than azithromycin. The two current anti-hypnozoite 8-aminoquinolines, primaquine(29) and tafenoquine(30, 31), require glucose-6-phosphate dehydrogenase deficiency testing which is an impediment (time, extra clinical visit, cost and accuracy of point of care test) to widespread drug implementation(32–35). The target indication for cethromycin is prevention of dormant *P. vivax* malaria after curative chloroquine or artemisinin combination treatment for primary *P. vivax* parasitemia. The INSPECTOR trial noted antagonism of tafenoquine with dihydroartemesinin-piperaquine for dormant *P. vivax*(36), which might be avoided with cethromycin’s different mechanism of action. Thus, cethromycin might be superior to daily doxycycline prophylaxis with shorter duration after malaria endemic country exit. Advantages to cethromycin are single daily doses, high liver tissue concentrations and broad antimicrobial activity. Next steps or validation of efficacy for dormant malaria in the *P. cynomolgi* model and good manufacturing practice and investigational new drug enabling preclinical studies for human malaria testing.

## MATERIALS AND METHODS

### Animals and parasites

Inbred, female 6-7-week-old wild-type BALB/c and C57Bl/6 mice were purchased from The Jackson Laboratory (Bar Harbor, ME), and kept in the animal facility at the Johns Hopkins Bloomberg School of Public Health. All animal experiments were approved by the Johns Hopkins Animal Care and Use Committee (ACUC), under protocol MO15H319. Sporozoite infected mCherry *P. berghei* mosquitoes were used for liver stage infections (37). *P. berghei* MRA-868 ANKA (*Pb*-ANKA-Luc) transgenic strain was used for all murine blood stage infections. The *Pb*-ANKA-Luc parasite strain expresses a GFP-luciferase fusion gene [19]. NF54 *P. falciparum* parasites at 4% hematocrit were cultured as described(38).

### Drug preparation and dosing

Primaquine, quinoline, and erythromycin were from Sigma. Azithromycin (Astatech cat # N460720) and cethromycin (BDG Synthesis, lot 16822, Wellington, New Zealand) were solubilized in either DMSO for in vitro assays or PBS/10% EtOH/20 mM HCl (875 μL PBS + 100 μL ethanol + 50 μL 1 N HCl per mL) for oral dosing in mice. For all the experimental studies done, each drug solution was dosed at a volume of 200 µL per mouse.

### Drug Measurements

Cethromycin concentrations in plasma and tissue were determined via liquid chromatography-tandem mass spectrometry by the Clinical Pharmacology Analytical Laboratory at Johns Hopkins. Briefly, cethromycin was isolated from plasma and methanol-homogenized tissue lysate via protein precipitation using a 0.45 µm Captiva filter plate (Agilent Technologies, Wilmington, DE). Cethromycin (C_42_H_59_N_3_O_10_) peak area ratios were normalized to the structural analog telithromycin (C_43_H_65_N_5_O_11_), which served as an internal standard. Cethromycin and its internal standard were quantified on an API 4000 (plasma) or API 4500 (tissue homogenates) (SCIEX, Foster City, CA) operated in positive ionization and selective reaction monitoring modes. Ion transitions monitored were *m/z* 766.5➔158.2 for cethromycin and *m/z* 813.6➔656.2 for telithromycin. The primary analytical measuring ranges for plasma and tissue assays were 0.1-2500 ng/mL and 10-2500 ng/sample, respectively. Tissue results were normalized to the weight of tissue analyzed. Assays were validated in accordance with regulatory recommendations.

### Liver stage in vitro Assay

Hepa 1-6 (ATCC CRL-1830) were grown to confluence in DMEM with Fungizone/10% Pencillin/Streptomycin and plated onto Permanox LAB-TEK slides. Approximately 50,000 *P. berghei* sporozoites were added per well. After 2 hours medium was washed with fresh medium containing drugs at 20 microM. At 24 hours after sporozoite infection, cells were washed with fresh medium alone or fresh medium with drugs. At 48 hours medium was removed and cells fixed with 4% paraformaldehyde for 15 minutes, washed 3 times with PBS for GFP-luciferase assays or in cold methanol for 15 minutes, dried and stored at -20C overnight for immunofluorescence. Antibodies were to mouse anti-*P. berghei* HSP70 (2E6) provided by Fidel Zavala and rabbit anti-Pb acyl carrier protein (ACP) provided by Dr S Prigge (Johns Hopkins University). Secondary FITC goat anti-mouse and Alexa594 goat anti-rabbit. Liver stage assay experiment was performed in biologic duplicate at two different dates.

### Model of sporozoite infection

Female *Anopheles stephensi* mosquitoes were fed with *mcherry P. berghei*-infected blood for sporozoite development, as previously described(37, 39). Twenty days later sort cage of 200 mosquitoes with fluorescent microscope, Infect mice with 5-7 mosquitoes, examine patency by Giemsa blood film microscopy starting from day 4 to day 16 post infection.

### Plasmodium falciparum culture

Low passage 3D7 or NF54 *Plasmodium falciparum* parasites were cultured and synchronized as described(40). Cultures were incubated with RPMI-1640 with L-glutamine, 25 mM HEPES, 40 µM hypoxanthine, 25 mM sodium bicarbonate, and 10% O+ human serum (Interstate Blood Bank). All cultures, unless otherwise noted, are in 2% hematocrit with O+ red blood cells. Cultures were gassed with 5% O_2_/ 5% CO_2_ /90% Nitrogen and placed in 37°C incubators.

### *In vivo* cytocidal model of murine malaria

The rodent model used for all the experiments were Balb/cj female mice from the Jackson Laboratories aged at least 6 weeks old and weighed approximately 20 grams each. For each study done, replicate of three mice were used for each drug dose regimen tested. *P. berghei* ANKA, 676m1cl1, Green fluorescence protein-luciferase (PbANKA GFP-Luc) obtained from ATCC (catalog # MRA-868) constitutively expresses the luciferase at all stages in the life cycle. For each experiment Balb/cj female mice were infected via intraperitoneal (i.p) with approximately 100,000 erythrocytes infected with PbANKA GFP-Luc from donor mouse between 5 to 10 % parasitemia (in first or second passage). For malaria blood stage infections and drug responses, there are not sex differences in Balb/c mice. As male and female Balb/c mice display differences in tissue iron levels, female mice were used for consistency throughout all experiments(41).

### Blood stage murine luciferase assay and analysis

During the drug treatment, 5 µL of blood was collected from the tail of each mouse at regular intervals and deposited into 45 µL of lysis buffer in a 96 well plate (29). Samples were stored at -80°C until processed. A total of 5 µL of blood/lysis buffer (whole blood equivalent of 0.5 μL) was transferred to a black, opaque 96-well plate and 95 µl of luciferase buffer (20 mM Tricine, 100 µM EDTA, 1.07 mM K_2_CO_3_, 2.67 mM MgSO_4_, 17 mM DTT, 250 µM ATP, 250 µM D-Luciferin) was added. Luciferase activity was measured in the IVIS Spectrum In Vivo Imaging System and analyzed using Living Image v. 4.4 software. The raw luciferase activity is expressed as radiant flux in photons/second. Total radiant flux was compared to parasites per well using GraphPad Prism 5 software

### Mosquito transmission

Gametocytes were cultured as previously described(42). Gametocytes (mature stages IV-V) were counted by blood film at day 15 to 16. Gametocyte cultures were then incubated with fresh medium alone or fresh medium with 10 μM drug for 48 hours, with medium replaced (with or without drug) after 24 hours exposure. Cultures were washed three times with fresh medium, and then diluted withRBCs and O+ human serum to a gametocytemia of 0.05-0.07%, 50% hematocrit, and kept warm at 37°C. Cultures were fed to 4-6 hour starved *Anopheles stephensi,* using the standard membrane feeding assay(42). After feeding, blood-fed mosquitoes were sorted and kept. Mosquitoes were maintained for 8 days, fed with 10% sucrose solution. At day 8, mosquitoes were dissected and midguts were stained with 1% mercurochrome to count oocysts.

### Pharmacokinetics

Pharmacokinetic (PK) data were collected at multiple time points from mice dosed orally with cethromycin at 30, 60, or 120 mg/kg from plasma, lung, and liver. For cross-tissue comparisons tissue concentrations in ng/mg were converted to ng/mL by multiplying by the approximate tissue density (lung ∼ 1000 mg/mL; liver ∼ 1050 mg/mL). Noncompartmental analyses were performed using the *PKNCA* (version 0.11.0) package in R (version 4.4.1). Mean tissue concentration-time profiles were generated for each dose group and tissue, and PKNCA was used to compute C_max_ and area under the concentration-time curve (AUC_0-72_) via the linear-up/log-down trapezoidal method. To estimate variability, bootstrapped NCA analyses (n=1000 resamples) were performed for each tissue-dose combination. To capture active elimination kinetics, and given that concentration-time profiles showed two phases—a rapid initial decline over 24 hours (alpha half-life, t_1/2α_) and a very slow terminal phase >100 hours, likely reflecting tissue redistribution—we calculated t_1/2α_ by fitting a linear regression of the log-transformed data for the first 24 hours.

## ACKNOWLEDGMENTS

We thank the Johns Hopkins Malaria Research Institute and The Bloomberg Family Foundation (D.J.S). This work was supported by NCATS R21 TR001737 titled “Quantum model repurposing of cethromycin for liver stage malaria” (DJS) and Maryland TEDCO funding (D.J.S. and N.K.). M.M.I. was supported by NIAID K23 AI139343 titled “Clinical pharmacology of malaria elimination” and by the Johns Hopkins Malaria Research Institute and The Bloomberg Family Foundation. We also thank Pamela Hummert, MS, in support of the drug concentration analyses.

## Author contributions

G.K. and D.S. designed experiments. M.M. designed and implemented pharmacokinetic results. M. M. and M.I. analyzed PK data. G.K., R.W. and K.P. implemented immunofluorescent hepatic assays. G.K. and B.B implemented liver stage transmission assays. R. W. implemented *P. falciparum* drug inhibition assays. G.K., B.B. and D.S. implemented murine blood stage assays. R.W., N.K. and D.S. implemented mosquito transmission assays. All authors analyzed the data. D.S. drafted manuscript. All authors provided critical review.

