## Appendix for "Cethromycin Pharmacokinetics and Pharmacodynamics for Single Dose Cure of *Plasmodium berghei* Liver Stages"

**Pharmacokinetics**

**Intra-assay precision, accuracy, dilutional integrity**

During the initial method development experiments, cethromycin and an isotopically-labeled internal standard, cethromycin-d6, were evaluated for selectivity, carryover, and cross talk. Although there were no interfering peaks observed in the six blank K_2_EDTA plasma lots or the three blank lots of tissue lysate at which cethromycin eluted (0.38 min in plasma and 1.05 min in tissue lysate; **supplement** **Figure 1**) and carryover was not observed, there was significant cross talk between cethromycin and the isotopically-labeled cethromycin-d6 (**Supplement Figure 2**). Consequently, the structural analog telithromycin was evaluated as an alternate internal standard for the quantification of cethromycin in plasma and tissue lysate. The use of the structural analog mitigated cross talk, and the molecule co-eluted with cethromycin (**supplement** **Figure 1**). The matrix effects determined via post column infusion demonstrated minimal ion suppression in plasma for cethromycin; conversely, in tissue lysate, both cethromcyin and telithromycin exhibited two- and four-fold ion enhancement, respectively.

Intra-assay precision and accuracy results for cethromycin in plasma and tissue lysate were as follows: plasma intra-precision ranged from 11.4% to 15.7% and inter-precision was 9.32% to 16.7%. In tissue lysate the intra- and inter-precision ranges were 6.52% to 12.1% and 3.70% to 10.9%, respectively. The intra- and inter-assay accuracy for cethromycin in plasma ranged from -3.12% to 5.67% and -8.58% to 11.6%; in tissue lysate the ranges were -5.06% to 11.2% and -11.5% to 16.3% respectively. A full summary of precision and accuracy studies is illustrated in **Table 1**.

The stability challenges included three freeze/thaw cycles at -20˚C and at -80˚C, unprocessed plasma and tissue lysate samples left at ambient temperature and protected from light for 24 hours, and finally, reconstituted samples left in the autosampler at 4˚C for 24 hours. The stability challenges were performed in quadruplicate at the low and high QC levels. The results of the stability studies met the aforementioned FDA bioanalytical guidelines of ≤15% difference of the unchallenged QC samples to the challenged QC samples.

The dilutional integrity studies for plasma samples diluted four- and 16-fold yielded %CVs ranging from 0.674% to 9.44% and percent accuracies ranging from 5.45% to 11.1% demonstrating that plasma samples may be diluted for analytical analysis. Tissue lysate samples demonstrated adequate extended linearity as well with %CVs ranging from 2.48% to 5.32% and percent accuracies ranging from 6.80% to 8.60% for the 64- and 256-fold diluted tissue lysate samples. The two-and four-fold dilutions performed on mid and high QC samples for evaluation of volume-limited samples generated precision and accuracy ranges from 9.32% to 14.8% and -10.5% to 7.50% in plasma; in tissue lysate the ranges were 3.22% to 8.97% and 7.50% to 13.6%, respectively (**Table 2**).

**Supplemental Figure 1 .** Representative chromatograms of (A) Cethromycin in plasma [100 ng/mL] and (B) Telithromycin [250 ng/mL] in plasma, with retention time demarcated at 0.37 min. (C) Cethromycin in tissue lysate [125 ng/sample] and (D) Telithromycin in tissue lysate [250 ng/sample], with retention time demarcated at 1.05 min

**Supplemental Figure 2.** Representative chromatogram of cross talk from the isotopically-labeled cethromycin-d6; (A) Chromatogram of cethromycin at the LLOQ in plasma (1.00 ng/mL) (black line) overlaid with the signal intensity of a blank plasma sample spiked only with isotopically-labeled cethromycin-d6 (blue line) and (B) Representative chromatogram of cethromycin-d6.

Supplemental Figure 1.

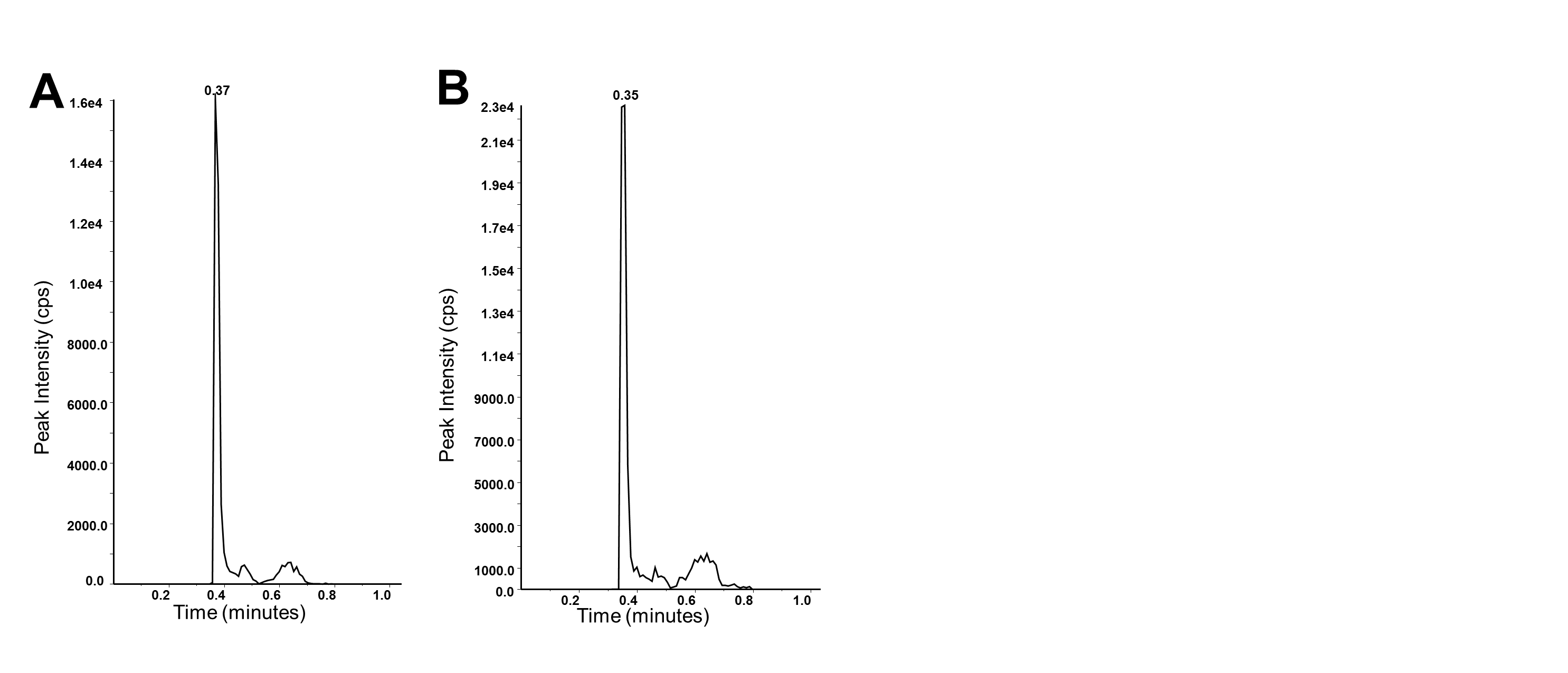

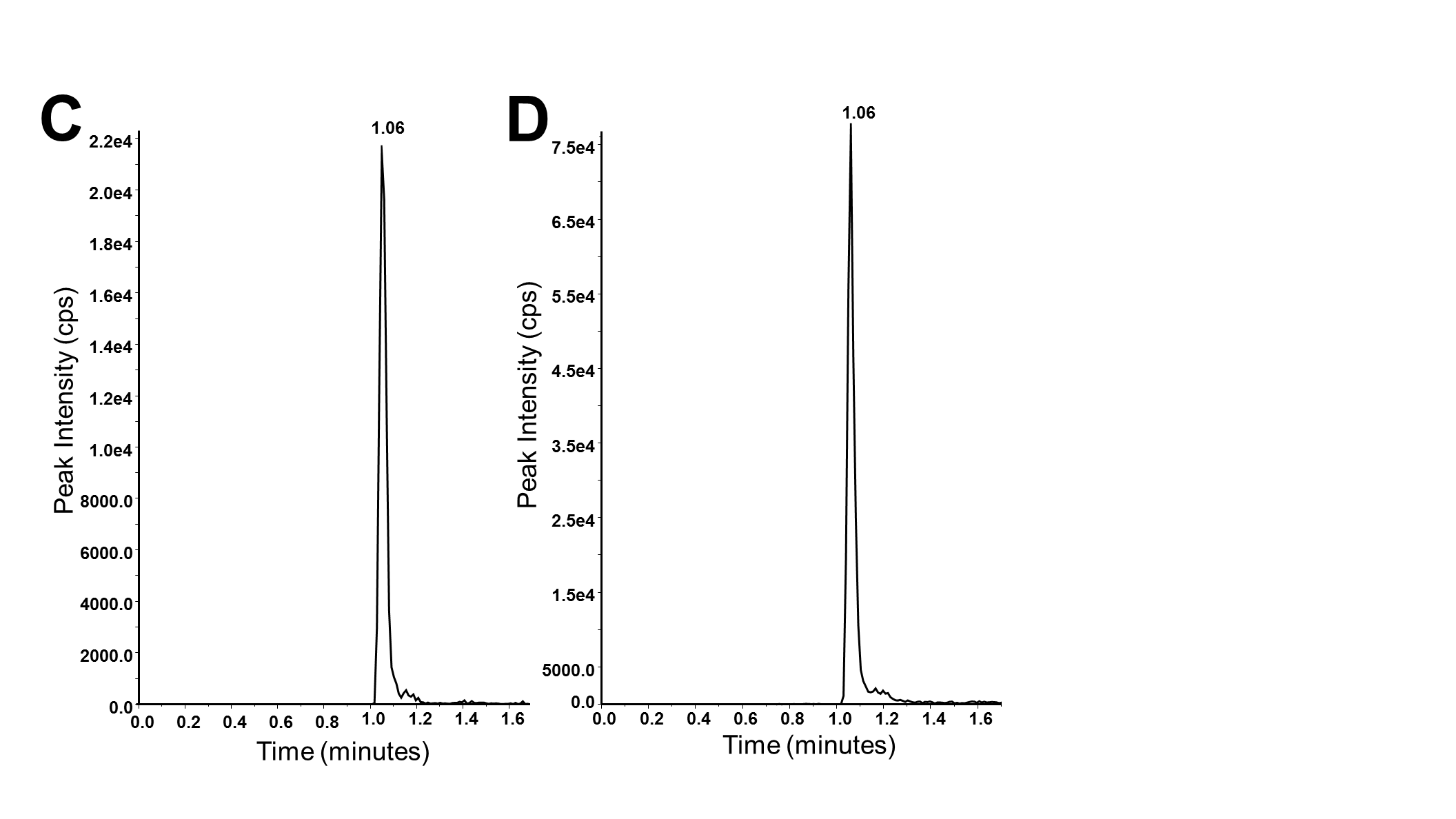

Supplemental Figure 2.

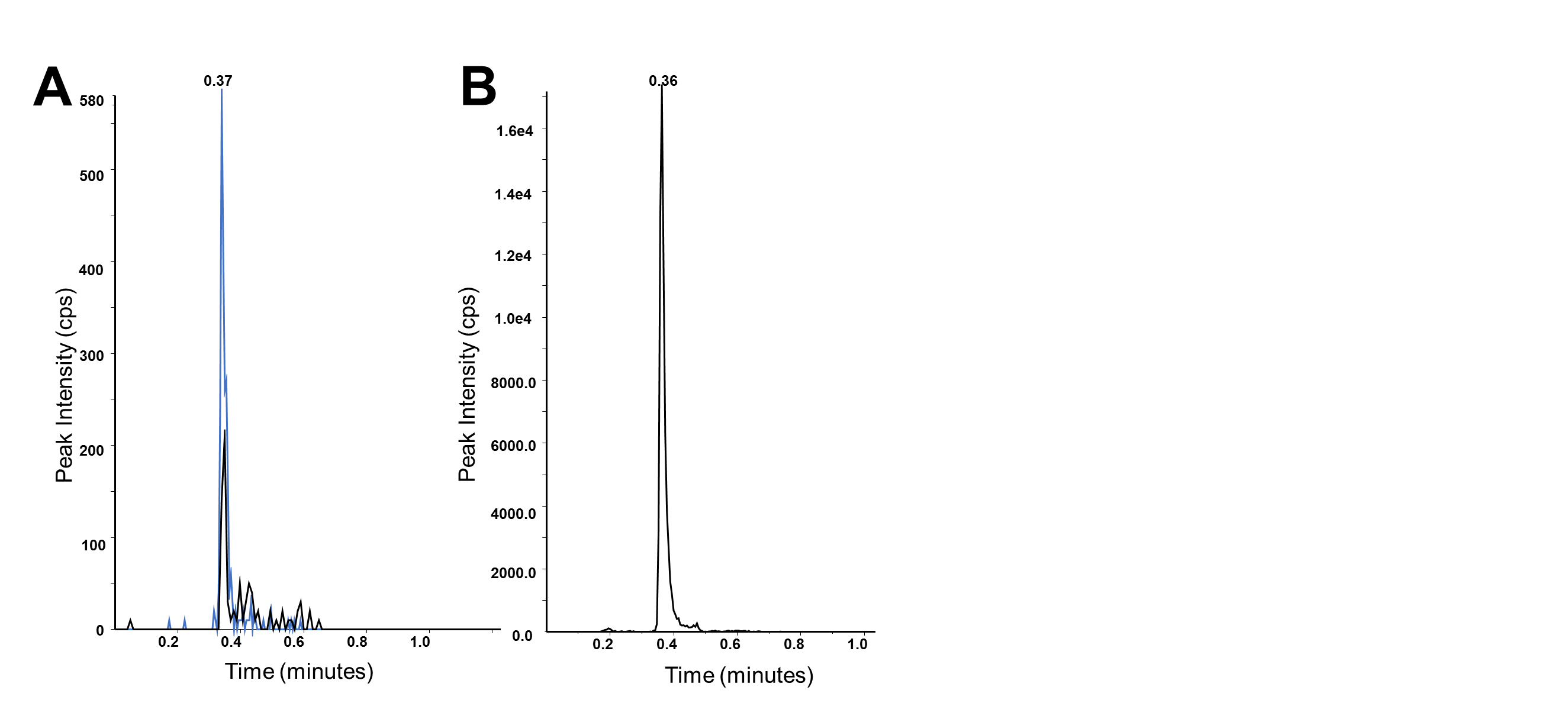

Table 1. Intra-assay and inter-assay precision and accuracy results.

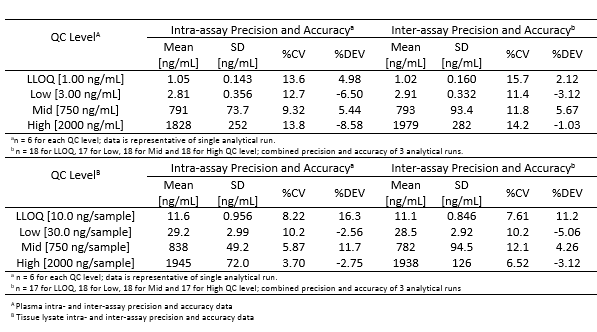

Table 2. Dilutional integrity sample studies in plasm and tissue lysate

| QC Level | [Mean] | [SD] | SD | %CV |
| --- | --- | --- | --- | --- |
| 3xULOQ [7500 ng/mL]  1/4 Dilution ^a^ | 7909 | 747 | 9.44 | 5.45 |
| 3xULOQ [7500 ng/mL]  1/16 Dilution ^b^ | 8331 | 56.2 | 0.674 | 11.1 |
| 50xULOQ [125000 ng/sample]  1/64 Dilution ^c^ | 135750 | 7228 | 5.32 | 8.60 |
| 50xULOQ [125000 ng/sample]  1/256 Dilution ^d^ | 133500 | 3317 | 2.48 | 6.80 |
| ^a^ n = 4 for the 1/4 dilution set in plasma  ^b^ n = 3 for the 1/16 dilution set in plasma | | | | |
| ^c^ n = 4 for the 1/64 dilution set in tissue lysate  ^d^ n = 4 for the 1/256 dilution set in tissue lysate | | | |  |
